# Peptidoglycan fragment release and NOD activation by commensal *Neisseria* species from humans and other animals

**DOI:** 10.1101/2024.01.09.574863

**Authors:** Tiffany N. Harris-Jones, Jia Mun Chan, Kathleen T. Hackett, Nathan J. Weyand, Ryan E. Schaub, Joseph P. Dillard

## Abstract

*Neisseria gonorrhoeae*, a human restricted pathogen, releases inflammatory peptidoglycan (PG) fragments that contribute to the pathophysiology of pelvic inflammatory disease. The genus *Neisseria* is also home to multiple species of human- or animal-associated *Neisseria* that form part of the normal microbiota. Here we characterized PG release from the human-associated nonpathogenic species *N. lactamica* and *N. mucosa* and animal-associated *Neisseria* from macaques and wild mice. An *N. mucosa* strain and an *N. lactamica* strain were found to release limited amounts of the pro-inflammatory monomeric PG fragments. However, a single amino acid difference in the PG fragment permease AmpG resulted in increased PG fragment release in a second *N. lactamica* strain examined. *Neisseria* isolated from macaques also showed significant release of PG monomers. The mouse colonizer *N. musculi* exhibited PG fragment release similar to that seen in *N. gonorrhoeae* with PG monomers being the predominant fragments released. All the human-associated species were able to stimulate NOD1 and NOD2 responses. *N. musculi* was a poor inducer of mouse NOD1, but *ldcA* mutation increased this response. The ability to genetically manipulate *N. musculi* and examine effects of different PG fragments or differing amounts of PG fragments during mouse colonization will lead to a better understanding of the roles of PG in *Neisseria* infections. Overall, we found that only some nonpathogenic *Neisseria* have diminished release of pro-inflammatory PG fragments, and there are differences even within a species as to types and amounts of PG fragments released.

## Introduction

The genus *Neisseria* contains multiple species of Gram-negative bacteria with varying cell morphology and niches. *Neisseria* species are almost always found associated with higher order host organisms, which includes multiple species of insects, birds and waterfowl, land and marine mammals, as well as the rhinoceros iguana and the marsupial quokka [1]. The most well-studied of the *Neisseria* species are *Neisseria gonorrhoeae* and *Neisseria meningitidis*, which are human-restricted pathogens and etiological agents for the sexually transmitted infection gonorrhea and invasive meningococcal disease, respectively. Most species in this genus are nonpathogenic colonizers of mucosal surfaces of their respective host organism. In fact, *N. meningitidis* has a nonpathogenic lifestyle as a colonizer of the naso- and oropharyngeal space of healthy adults [2]. Like *N. meningitidis*, the major niche occupied by commensal mammal-associated *Neisseria* is the oral and nasopharyngeal space. *Neisseria* are considered part of the core oral microbiome of healthy humans and constitute up to 8% of the oral microbiota of humans [3-5].

The nonpathogenic species of *Neisseria* used in this study are all colonizers of mammals. *Neisseria lactamica* and *Neisseria mucosa* are associated with humans. The newly recognized species, *Neisseria musculi*, was isolated from wild house mice [6]. The macaque isolates AP312 and AP678 were isolated from macaques housed in a national primate center [7]. In terms of cell morphology, *N. lactamica*, *N. mucosa*, and macaque isolates AP312 and AP678 are all diplococci, while *N. musculi* is a rod-shaped bacterium [1, 6, 7]. There are temporal and tissue-specific colonization patterns in different species of nonpathogenic *Neisseria*. For example, *N. lactamica* carriage rates are higher in children compared to adults, and nasopharyngeal colonization by *N. lactamica* protects against colonization by *N. meningitidis* [8-10]. In contrast, *N. meningitidis* is commonly found in the throat of adults [8, 11]. *N. mucosa* is mostly found in the gingival plaque and tooth surfaces of children and adults but does not appear to contribute to exacerbating or preventing tooth decay [11, 12]. *N. musculi* was isolated from the mouths of wild mice but is able to colonize both the oral cavity and the gastrointestinal tract of mice in the laboratory [13]. Although the macaque isolates colonize the nasopharyngeal and oral cavities of rhesus macaques, AP312 was initially isolated from a bite wound [7].

Despite their roles as part of the core oral microbiome, nonpathogenic human-associated *Neisseria* encode several virulence-associated factors and in rare cases will cause disease in people who are immunocompromised, have pre-existing risk factors, or post-trauma [14, 15]. Some diseases caused by commensal *Neisseria* spp. include high fatality diseases like meningitis, septicemia, and endocarditis, as well as conjunctivitis, respiratory tract infections and pneumonia [14]. Multiple cases of endocarditis by commensal *Neisseria* spp. have occurred after dental procedures most likely due to oral wounds sustained from these procedures that provided a direct route for the bacteria to travel from the mouth to the bloodstream [14]. Factors that favor colonization by these nonpathogenic *Neisseria* species while not promoting inflammation and disease are not clear.

One factor that was previously found to differ between the inflammation inducing species *N. gonorrhoeae* and the more commensal species *N. meningitidis*, *N. mucosa*, and *N. sicca* is the release of peptidoglycan fragments, with the commensal species releasing about one-third as much of the pro-inflammatory molecules [16]. Peptidoglycan (PG) makes up the bacterial cell wall and confers cell shape and protection against osmotic shock. PG consists of a glycan backbone of repeating subunits of *N*-acetylglucosamine (GlcNAc) and *N-*acetylmuramic acid (MurNAc), with peptide stems extending off MurNAc that can be crosslinked to adjacent peptide stems forming a mesh-like structure. Because of PG remodeling to allow for cell enlargement and cell separation, small PG fragments are liberated from the sacculus during growth [17]. In Gram-negative bacteria, these PG fragments are usually taken back into the cytoplasm to be recycled for reuse in the cell wall or general cellular metabolism. A limited number of Gram-negative bacteria, including the human pathogens *N. gonorrhoeae, N. meningitidis*, and *Bordetella pertussis* release sufficient amounts of pro-inflammatory PG fragments to stimulate production of pro-inflammatory cytokines in tissue explants [18-21]. NOD1 and NOD2 are immune receptors found in host cells that are activated by small PG fragments [22, 23].

In this study, we characterized the PG fragments released by nonpathogenic human- and animal-associated *Neisseria* and their ability to activate NOD1 and/or NOD2. We hypothesized that lower amounts of PG fragment release might allow for continued colonization and not produce inflammatory responses that could lead to clearance of the bacteria. We found that PG fragment release from human-associated species *N. mucosa* and one strain of *N. lactamica* was similar to that previously seen in *N. meningitidis* or *N. sicca*, with low amounts of PG monomer released and more of the PG fragments being broken down into free sugars and free peptides. However, a different strain of *N. lactamica* exhibited PG fragment release similar to *N. gonorrhoeae* with large amounts of PG monomers released. We found that an amino acid difference in the PG fragment permease AmpG was responsible for the differing amounts of PG fragments released by *N. lactamica*, and a minority of strains encode the inefficient AmpG variant. The animal-associated species also released PG fragments, with some differences in the amounts of specific types of fragments released. *N. musculi* showed a PG fragment profile similar to *N. gonorrhoeae*, though with lower amounts of PG released. Mutations affecting PG processing or recycling in *N. musculi* were made and affected amounts of PG released and mouse NOD1-mediated responses.

## Materials and methods

### Bacterial strains and growth conditions

All strains used are listed in Table 1. All *Neisseria* strains are grown at 37°C either on gonococcal base medium (Difco) agar plates (GCB) with 5% CO2 or in gonococcal base liquid medium with 0.042% NaHCO3 and Kellogg’s supplements (cGCBL) [24, 25]. When necessary, 80 μg/ml kanamycin was added to the growth medium. *Escherichia coli* strains were grown on LB agar or in LB broth (Difco) at 37°C. The growth medium was supplemented with 40 μg/ml kanamycin, 500 μg/ml erythromycin, or 25 μg/ml chloramphenicol as needed.

**Table 1.**
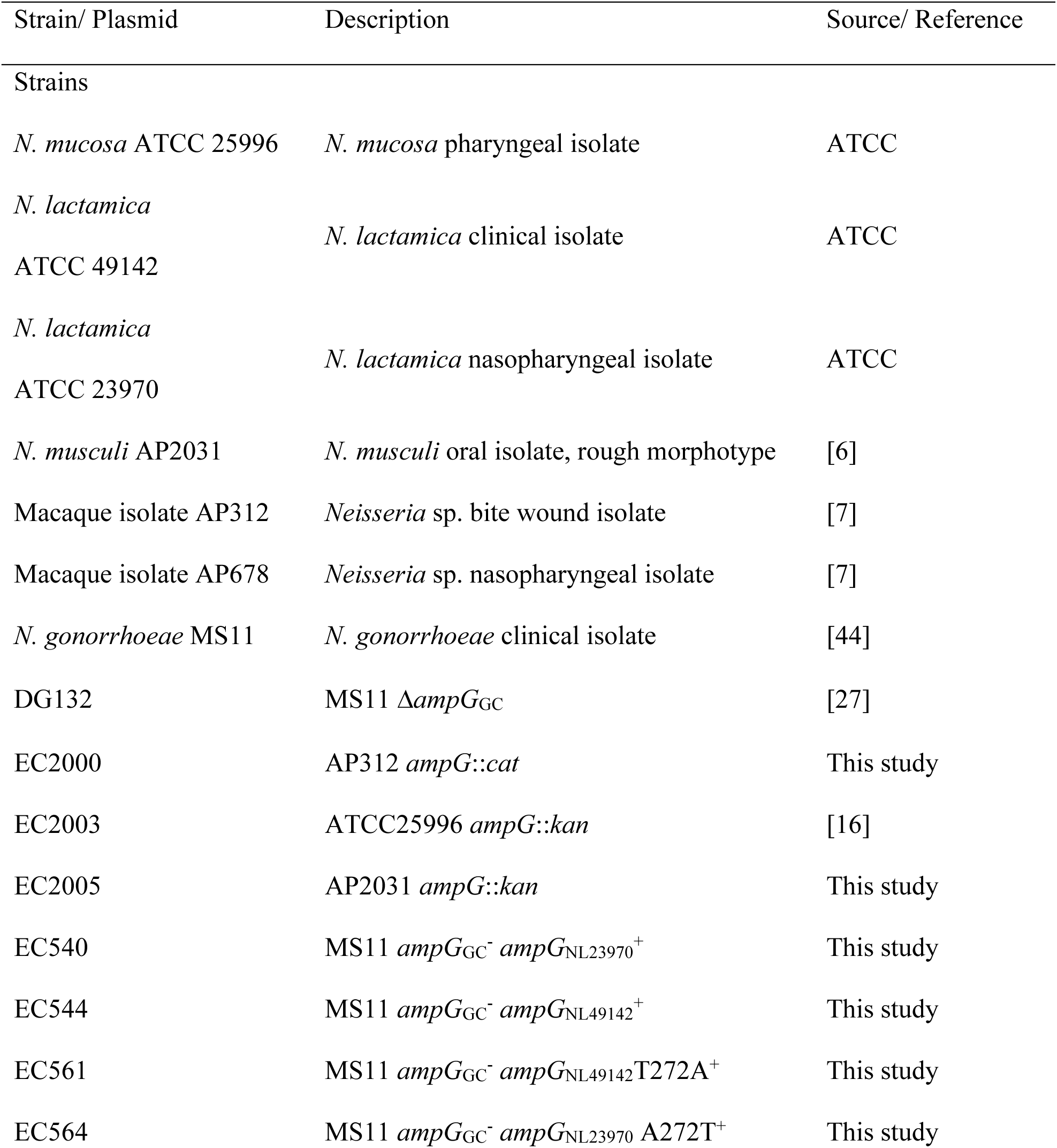
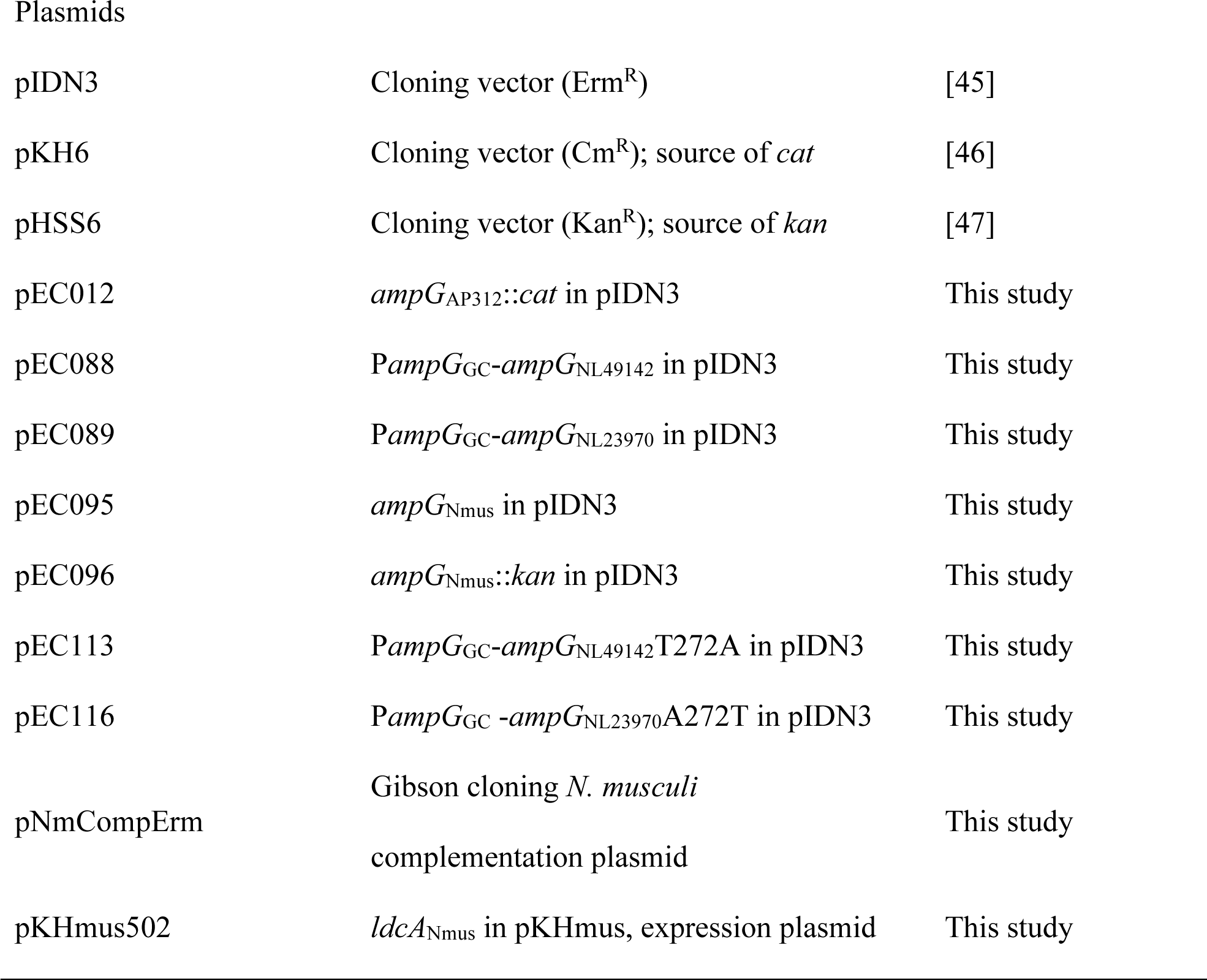
Strains and plasmids used in this study.

| Strain/ Plasmid | Description | Source/ Reference |
| --- | --- | --- |
| Strains |  |  |
| *N. mucosa* ATCC 25996 | *N. mucosa* pharyngeal isolate | ATCC |
| *N. lactamica*<br>ATCC 49142 | *N. lactamica* clinical isolate | ATCC |
| *N. lactamica*<br>ATCC 23970 | *N. lactamica* nasopharyngeal isolate | ATCC |
| *N. musculi* AP2031 | *N. musculi* oral isolate, rough morphotype | [6] |
| Macaque isolate AP312 | *Neisseria* sp. bite wound isolate | [7] |
| Macaque isolate AP678 | *Neisseria* sp. nasopharyngeal isolate | [7] |
| *N. gonorrhoeae* MS11 | *N. gonorrhoeae* clinical isolate | [44] |
| DG132 | MS11 $\Delta ampG_{GC}$ | [27] |
| EC2000 | AP312 *ampG::cat* | This study |
| EC2003 | ATCC25996 *ampG::kan* | [16] |
| EC2005 | AP2031 *ampG::kan* | This study |
| EC540 | MS11 *ampG<sub>GC</sub><sup>-</sup> ampG<sub>NL23970</sub><sup>+</sup>* | This study |
| EC544 | MS11 *ampG<sub>GC</sub><sup>-</sup> ampG<sub>NL49142</sub><sup>+</sup>* | This study |
| EC561 | MS11 *ampG<sub>GC</sub><sup>-</sup> ampG<sub>NL49142</sub>T272A<sup>+</sup>* | This study |
| EC564 | MS11 *ampG<sub>GC</sub><sup>-</sup> ampG<sub>NL23970</sub> A272T<sup>+</sup>* | This study |
| Plasmids |  |  |
| pIDN3 | Cloning vector (Erm<sup>R</sup>) | [45] |
| pKH6 | Cloning vector (Cm<sup>R</sup>); source of *cat* | [46] |
| pHSS6 | Cloning vector (Kan<sup>R</sup>); source of *kan* | [47] |
| pEC012 | *ampG*<sub>AP312</sub>::*cat* in pIDN3 | This study |
| pEC088 | *PampG*<sub>GC</sub>-*ampG*<sub>NL49142</sub> in pIDN3 | This study |
| pEC089 | *PampG*<sub>GC</sub>-*ampG*<sub>NL23970</sub> in pIDN3 | This study |
| pEC095 | *ampG*<sub>Nmus</sub> in pIDN3 | This study |
| pEC096 | *ampG*<sub>Nmus</sub>::*kan* in pIDN3 | This study |
| pEC113 | *PampG*<sub>GC</sub>-*ampG*<sub>NL49142</sub>T272A in pIDN3 | This study |
| pEC116 | *PampG*<sub>GC</sub> -*ampG*<sub>NL23970</sub>A272T in pIDN3 | This study |
| pNmCompErm | Gibson cloning *N. musculi*<br>complementation plasmid | This study |
| pKHmus502 | *ldcA*<sub>Nmus</sub> in pKHmus, expression plasmid | This study |

### Strain and plasmid construction

All strains and plasmids used in this study are listed in Table 1, and primers are listed in Table 2. Spot transformation was used to generate *Neisseria* mutants [26]. Briefly, around 0.5 - 1 μg plasmid DNA was digested with PciI with a subsequent heat-inactivation step to linearize the plasmid, and the digest reaction was spotted onto a pre-warmed GCB agar plate. Five to ten piliated colonies were then streaked over the DNA spots, and the plate was incubated at 37°C overnight. Colonies growing on the spots were restreaked onto fresh GCB plates for selection or screening. All transformants were screened by PCR and sequencing.

**Table 2.**
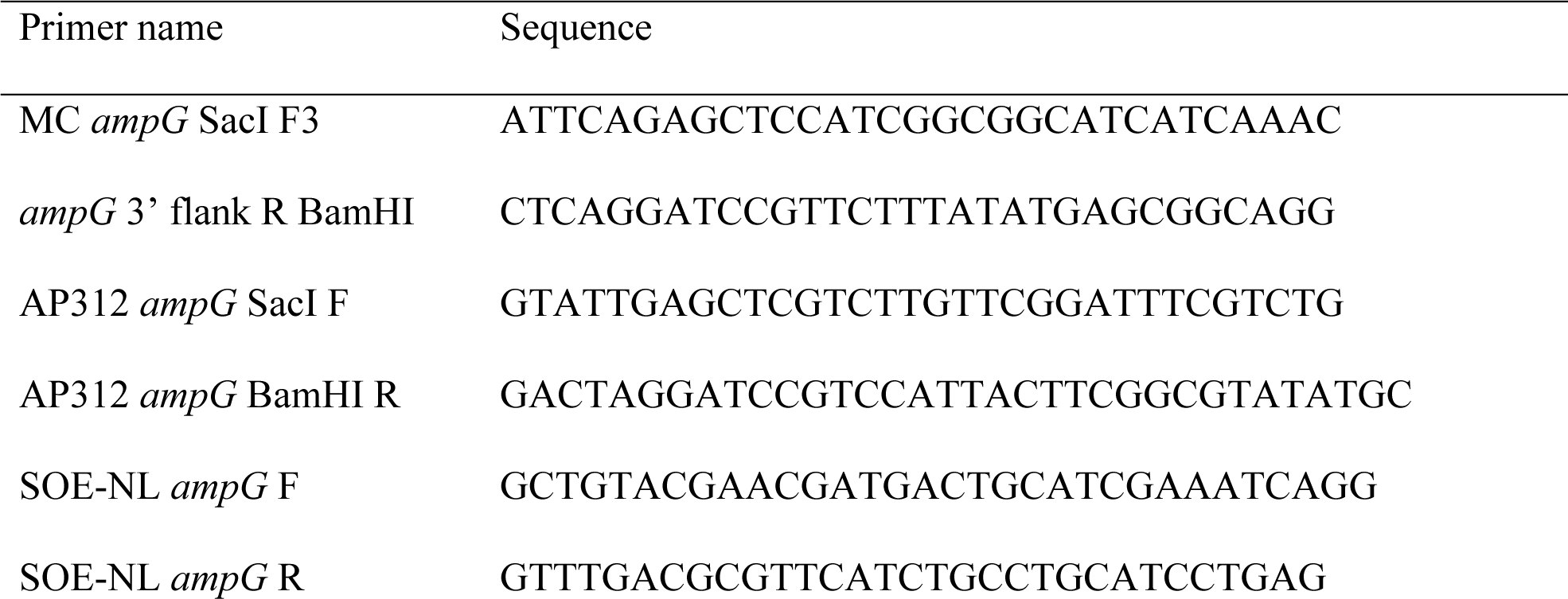
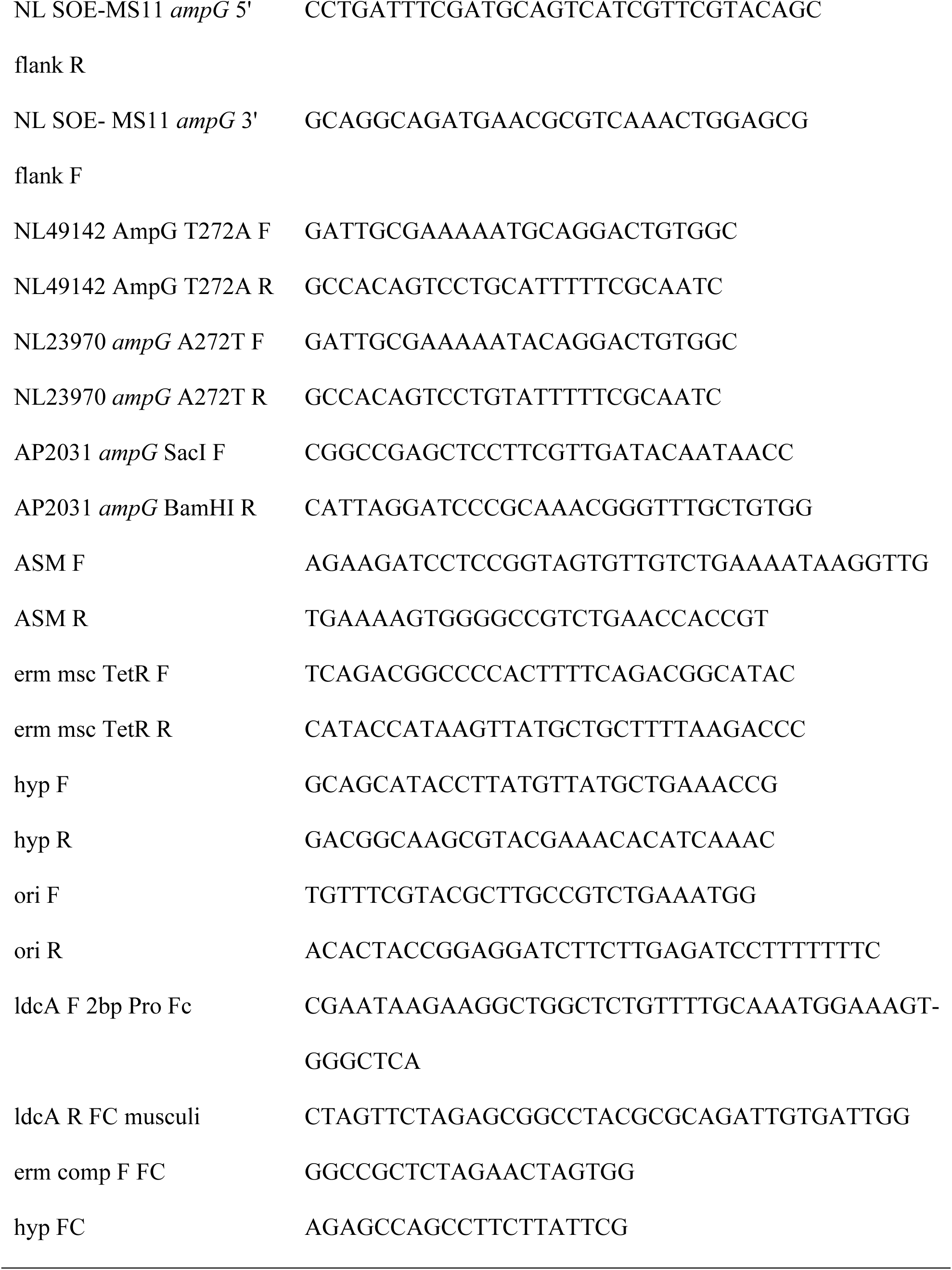
Primers used in this study.

To construct pEC012 (AP312 *ampG*::*cat* in pIDN3), AP312 *ampG* was first amplified from macaque isolate AP312 chromosomal DNA with primers AP312 *ampG* SacI F and AP312 *ampG* BamHI R, and subsequently digested with SacI, BamHI, DpnI and EarI, resulting in two DNA fragments. The chloramphenicol resistance gene *cat* was excised from pKH6 by digestion with DpnI and EarI. The cloning vector pIDN3 was digested with SacI and BamHI. All four digest products were ligated together to form pEC012. Macaque isolate AP312 was transformed with pEC012 to make EC2000.

The insert for pEC088 (P*ampG*GC-*ampG*NL49142 in pIDN3) was generated via overlap-extension-PCR (OE-PCR). First, *ampG*NL49142 was amplified from *N. lactamica* ATCC 49142 chromosomal DNA using primers SOE-NL *ampG* F and SOE-NL *ampG* R. Approximately 1kb *ampG*GC upstream and downstream regions, which include the native *ampG*GC promoter and transcriptional terminator, were amplified from *N. gonorrhoeae* MS11 chromosomal DNA using primer pairs MC *ampG* SacI F3/NL SOE-MS11 *ampG* 5’ flank R and NL SOE-MS11 *ampG* 3’ flank F/*ampG* 3’ flank R BamHI, respectively. The three PCR products were used as templates in OE-PCR with primes MC *ampG* SacI F3 and *ampG* 3’ flank R BamHI. The final PCR product was digested with SacI and BamHI and ligated into similarly digested pIDN3 to form pEC088. pEC089 (P*ampG*GC-*ampG*NL23970 in pIDN3) was constructed almost exactly as pEC088; the only exception is that *ampG*NL23970 was amplified from *N. lactamica* ATCC 23970 chromosomal DNA. Transformation of *N. gonorrhoeae* MS11 with pEC088 or pEC089 yielded EC544 and EC540, respectively.

To generate pEC095 (*ampG*Nmus in pIDN3), *ampG*Nmus was amplified from *N. musculi* AP2031 chromosomal DNA using primers AP2031 *ampG* SacI F and AP2031 *ampG* BamHI R, digested with SacI and BamHI and ligated into similarly digested pIDN3. Plasmid pEC095 was digested with BtsBI and the 5’ overhanging DNA ends were filled in with T4 Polymerase (NEB). A kanamycin resistance marker, *kan*, was excised from pHSS6 by digestion with NheI and BamHI, treated with T4 Polymerase and blunt-ligated with the BtsBI digested pEC095 to form pEC096 (*ampG*Nmus::*kan* in pIDN3). Plasmid pEC096 was transformed into *N. musculi* AP2031 to generate EC2005.

Plasmid pEC088 was used as template in two PCR reactions with primer pairs MC *ampG* SacI F3/NL49142 *ampG* T272A F and NL49142 *ampG* T272A R/*ampG* 3’ flank R BamHI. The primer sequences of NL49142 *ampG* T272A F and NL49142 *ampG* T272A R contain a missense mutation (ACA->GCA) that would result in substitution of residue 272 from a threonine to an alanine in the final gene product. The two PCR products were used in OE-PCR with primers MC *ampG* SacI F3 and *ampG* 3’ flank R BamHI, and the final OE-PCR product was digested with SacI and BamHI and ligated into similarly digested pIDN3 to form pEC113 (P*ampG*GC-*ampG*NL49142T272A in pIDN3). pEC116 (P*ampG*GC-*ampG*NL23970A272T in pIDN3) was built with a similar strategy as pEC113, with two variations during the initial PCR reaction. Chromosomal DNA from EC540 was used as template in the PCR reaction, with primer pairs MC *ampG* SacI F3/NL23970 *ampG* A272T F and NL23970 *ampG* A272T R/*ampG* 3’ flank R BamHI instead. *N. gonorrhoeae* MS11 was transformed with pEC113 or pEC116 to form EC561 and EC564, respectively.

To create a complementation plasmid for *N. musculi*, the adenine-specific methyltransferase and a downstream gene for a hypothetical protein were amplified from AP2031 (rough isolate) chromosomal DNA using primers ASM-F and ASM-R in addition to hyp-F and hyp-R. The plasmid origin was amplified from pIDN1 using oriF and oriR, and a region containing *ermC*, the *tet* repressor and inducible promoter/operator were amplified from pKH18 using primers erm mcs TetR F and erm mcs TetR R. The final plasmid pNmCompErm was constructed using Gibson assembly.

An *ldcA* overexpression plasmid for *N. musculi* was created by amplifying a constitutively-expressed version of *ldcA* using primers ldcA 2bp F Pro Fc and ldcA R FC musculi and amplifying the pNmCompErm plasmid using primers erm comp F FC and hyp FC. The final plasmid pKHmus2 was created using Gibson assembly.

### Metabolic labeling of peptidoglycan with [^3^H] glucosamine or [^3^H] DAP and quantitative fragment release

Metabolic labeling of PG with [6-^3^H]glucosamine or [2,6-^3^H]DAP and quantitative fragment release was performed as described previously for *N. gonorrhoeae* [16, 27]. For [6-^3^H]glucosamine labeling, strains were grown in cGCBL to mid-log phase, diluted to OD540 of 0.2 and pulse labeled with 10 μCi/ml [6-^3^H]glucosamine in GCBL supplemented with 0.042% NaHCO3 and modified Kellogg’s supplements containing pyruvate instead of glucose for 30 minutes. The cells were washed to remove unincorporated label and resuspended in GCBL containing supplements and glucose for the chase period. The cells were grown for 2.5 hours, after which cell-free supernatant was harvested by centrifugation at 3,000 x g for 10 minutes and filtration using a 0.22 μm filter. For quantitative fragment release, the amount of radiation (counts per minute, CPM) in the cell pellets were determined by liquid scintillation counting prior to the chase period and normalized to each other. Labeling with [2,6-^3^H]DAP was performed in a similar way as labeling with [6-^3^H]glucosamine, with variations in the pulse labeling phase. Strains were grown in DMEM lacking cysteine supplemented with 25 μCi/ml [2,6-^3^H] DAP, 100 μg/ml threonine and 100 μg/ml methionine for 60 minutes for the labeling phase. Released radiolabeled PG fragments are separated by size-exclusion chromatography and detected by liquid scintillation counting.

### NOD1 and NOD2 activation with HEK293-reporter cells

NOD1 and NOD2 activation were measured using HEK293-reporter cells. Briefly, bacterial strains were grown to mid-log phase in cGCBL from an initial OD540 of 0.2. Cells were removed by centrifugation, and supernatants were passed through a 0.2 µM filter. Supernatants were normalized by dilution with GCBL, based on total protein content of cells in the cultures, and added to HEK293- reporter cells overexpressing NOD1 or NOD2 as described in (29). NF-kB activation was measured at OD650 after incubation of cell supernatants in QUANTI-Blue^TM^ medium as described in manufacturer’s instructions (InvivoGen).

## Results

### Nonpathogenic *Neisseria* release PG fragments

PG fragments released by growing *N. gonorrhoeae* and *N. meningitidis* have been previously characterized through the method of metabolic pulse-chase labeling of the PG with [^3^H]-glucosamine (glcNH2) and [^3^H]-diaminopimelic acid (DAP), which labels the glycan backbone and peptide stems of PG, respectively [19, 28-30]. [^3^H]-glucosamine labeled PG fragments released by WT cells are PG dimers, tetrasaccharide-peptide, PG monomers, free disaccharide, and free anhydro-*N*-acetylmuramic acid (anhMurNAc). [^3^H]-DAP labeled PG fragments are PG dimers, tetrasaccharide-peptide, PG monomers, free tetrapeptide and free tripeptide (in one peak), and free dipeptide (in a second peptide peak). Any DAP that was unincorporated will appear as a shoulder on the dipeptide peak. The known structures of the released PG fragments are summarized in Fig. 1A. The major inflammatory molecules released are the PG dimers, PG monomers, and free tetra- and tri-peptide (Fig. 1B) [29]. PG dimers that are digested by host lysozyme yield PG monomers with reducing ends, and these molecules are NOD2 agonists [31]. The tripeptide PG monomers and the free tripeptides are human NOD1 agonists, while the tetrapeptide PG monomers and free tetrapeptides are mouse NOD1 agonists [22, 32]. Small amounts of dipeptide PG monomers are released and serve as NOD2 agonists [33].

**Figure 1.**
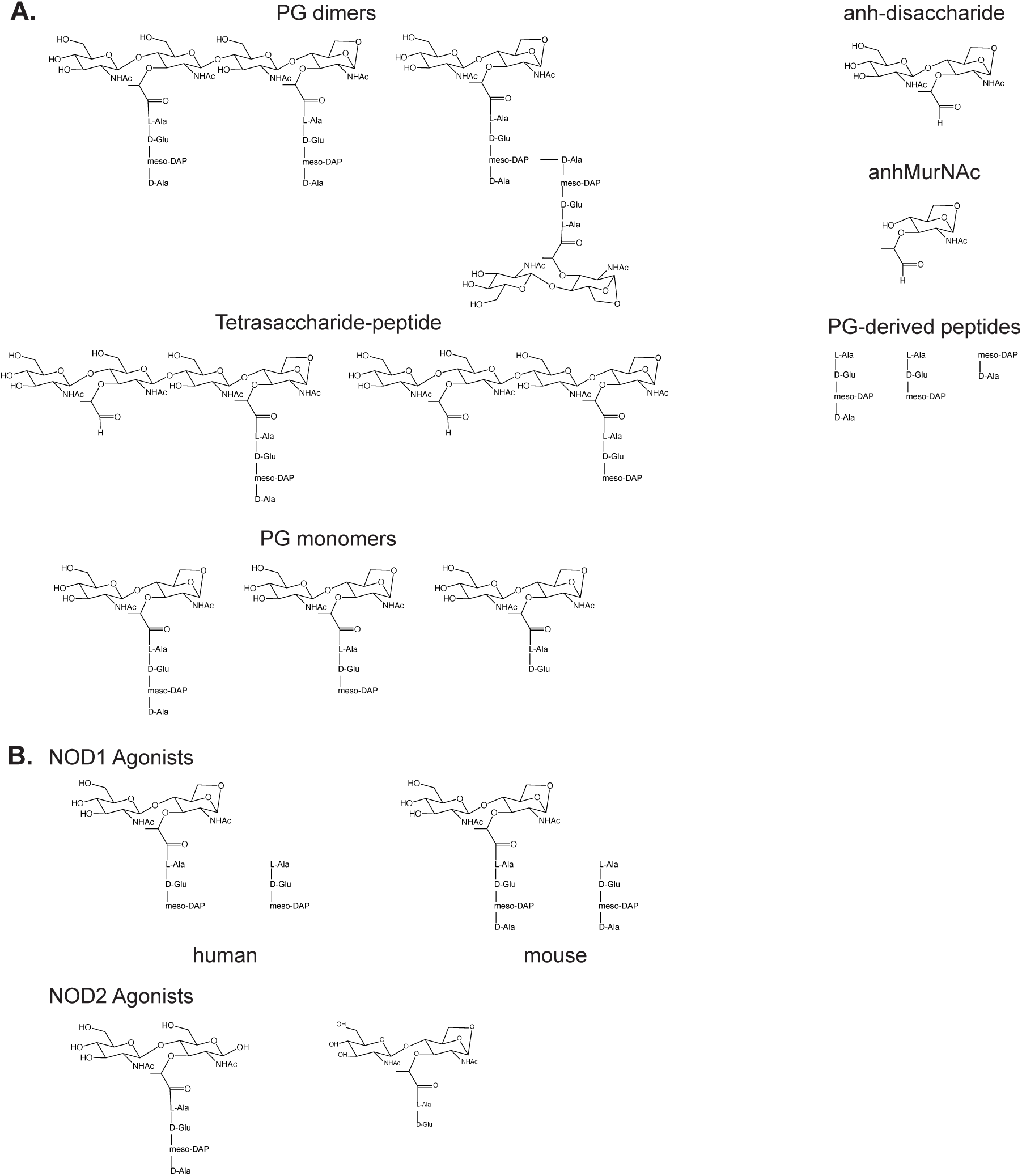
A. Peptidoglycan fragments released into the milieu by *Neisseria* species. B. Peptidoglycan fragments that stimulate the human or mouse pattern recognition receptor NOD1, or that stimulate NOD2 in both species.

We performed metabolic, pulse-chase labeling of the PG of the human-associated species *N. mucosa*. Consistent with previous studies, *N. mucosa* labeled with [6-^3^H]-glucosamine released PG monomers, free disaccharide, and anhydro-MurNAc but did not release any detectable PG dimers (Fig. 2A). To also examine PG-derived peptides released by *N. mucosa*, we used metabolic labeling with [2,6-^3^H]-DAP. DAP-labeled PG fragments released by *N. mucosa* included PG monomers, free tetrapeptide and tripeptide, and free dipeptide (Fig. 2B). Again, no PG dimer release was detected. A quantitative comparison with PG fragments released by *N. gonorrhoeae* showed that *N. mucosa* released lower amounts of PG monomer, as previously noted [16], and released larger amounts of free tetrapeptide and tripeptide (Fig. 2B). Thus PG monomer release in *N. mucosa* is mostly similar to what is seen with *N. meningitidis*, with larger amounts of PG fragments being broken down to free peptides and sugars and less being released as intact PG monomers.

**Figure 2.**
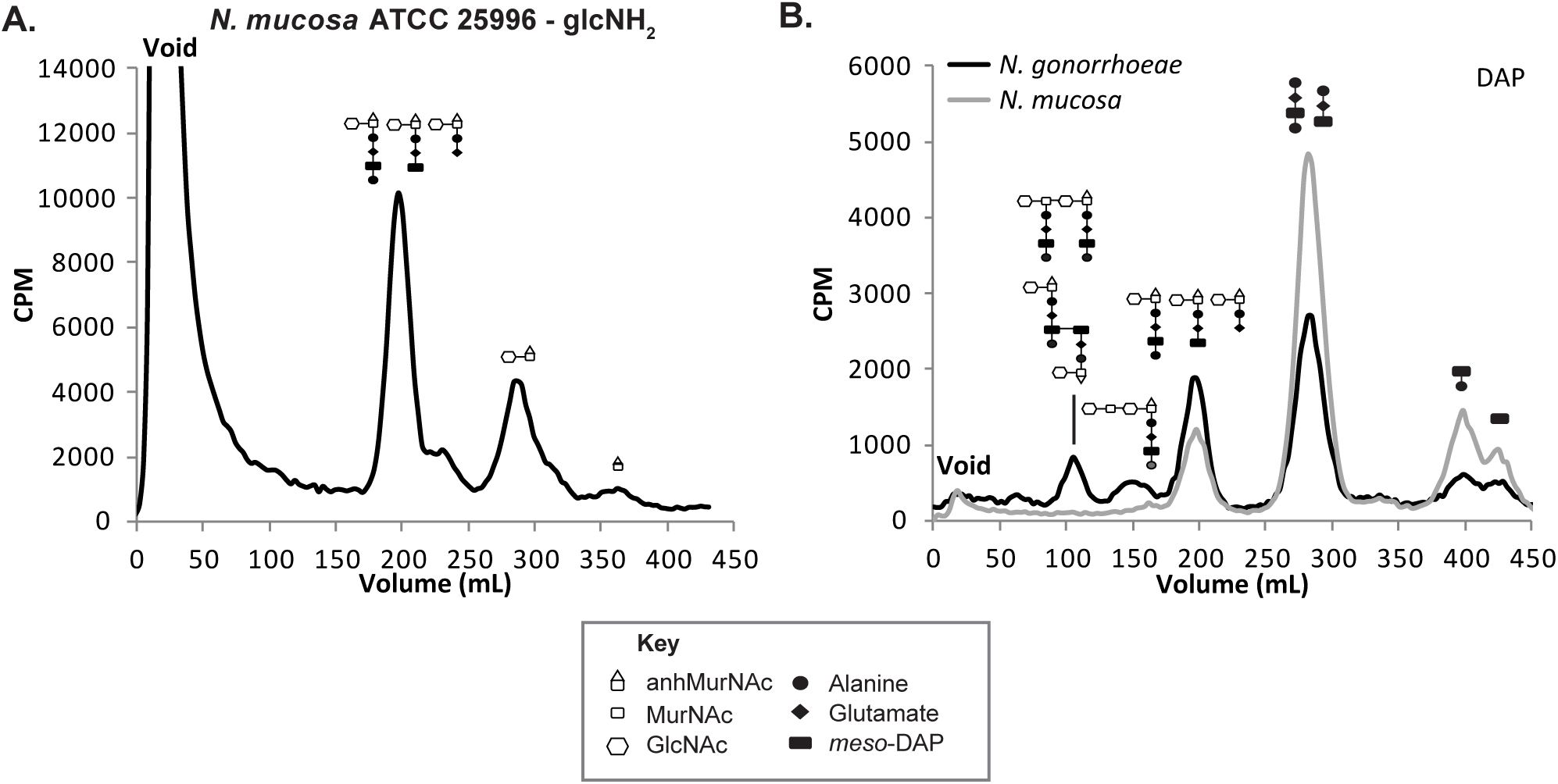
Nonpathogenic human-associated species *Neisseria mucosa* strain ATCC 25996 releases PG fragments into the medium, and the PG fragments released are separated by size-exclusion chromatography. A. PG fragments released during the chase period following metabolic labeling of PG with [^3^H]-glucosamine (glcNH2). B. PG fragments released from *N. mucosa* following [^3^H]-DAP labeling compared quantitatively to those released by *N. gonorrhoeae*. Cartoon depictions of the PG fragments found in gonococcal supernatants use symbols from Jacobs et al. [34].

We examined PG fragment release in the human-associated species *N. lactamica,* examining strains ATCC 23970 and ATCC 49142. Attempts to metabolically label PG in these two isolates using [6-^3^H]-glucosamine were not successful. Therefore, we turned to labeling of the PG peptide chains using [2,6-^3^H]-DAP. The PG fragment release profile of ATCC 23970 showed characteristics similar to *N. mucosa* in that ATCC 23970 released larger amounts of free peptides and lower amounts of PG monomers (Fig. 3A). Unlike *N. mucosa*, ATCC 23970 released significant amounts of PG dimers and tetrasaccharide-peptide. Overall, these data indicate that *N. lactamica* ATCC 23970 is similar to *N. mucosa* and *N. meningitidis* in substantially breaking down PG fragments via the AmiC-LtgC pathway prior to fragment release [30]. By contrast, *N. lactamica* ATCC 49142 PG fragments released were made up of PG monomers and free peptides in approximately equal amounts, similar to PG fragment release seen with *N. gonorrhoeae* (Fig. 3B). Like ATCC 23970, ATCC 49142 also released PG dimers.

**Figure 3.**
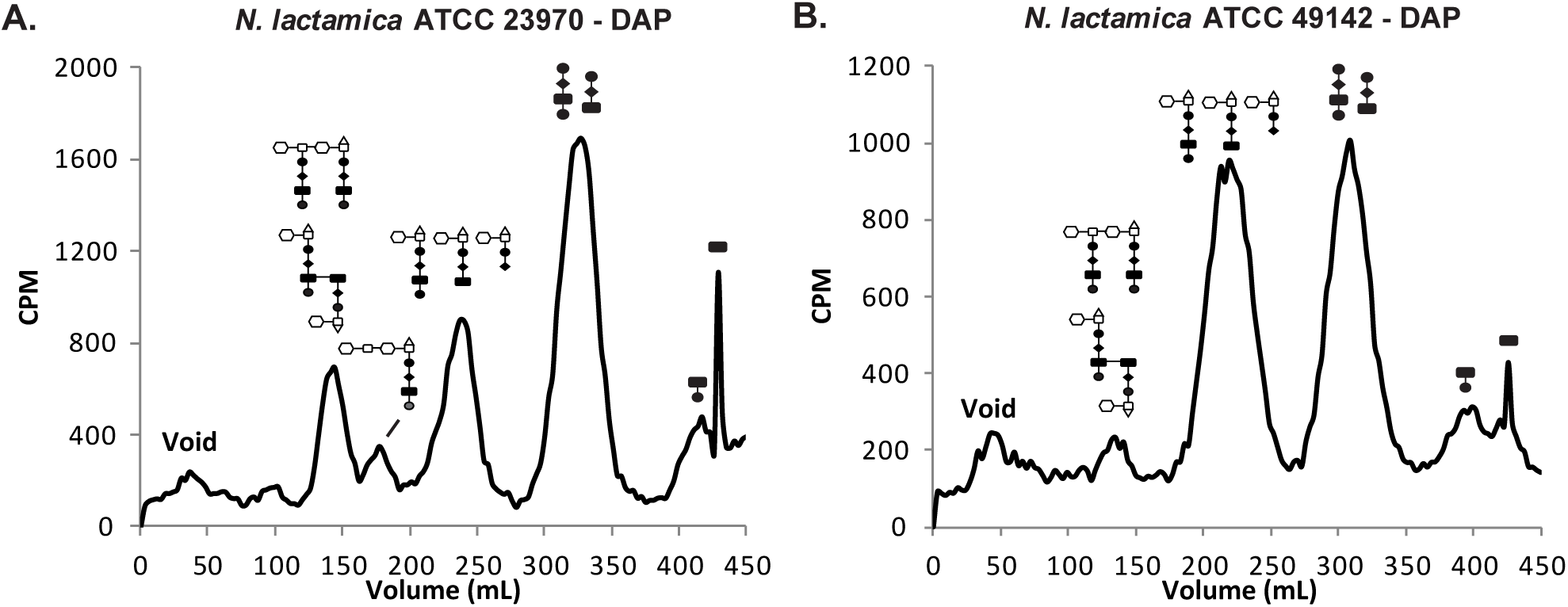
PG fragments released by *Neisseria lactamica* strain ATCC23970 (A) or *N. lactamica* strain 49142 (B) following metabolic labeling with [^3^H]-DAP.

### The difference in PG monomer release by *N. lactamica* ATCC 23970 and *N. lactamica* ATCC 49142 is partially due to a single nucleotide polymorphism in AmpG

A major difference in PG fragment release between *N. gonorrhoeae* and *N. meningitidis* is due to amino acid differences in the sequence of the PG fragment permease AmpG [16],. AmpG functions in both species to transport anhydro-disaccharide containing PG fragments into the cytoplasm for recycling, but due to the amino acid differences, *N. gonorrhoeae* AmpG is less efficient and 3-4 fold more PG monomers are released by *N. gonorrhoeae* than by *N. meningitidis*.

To examine PG recycling in *N. lactamica*, we attempted to mutate *N. lactamica ampG*, but we were unable to genetically manipulate either of the *N. lactamica* isolates. As an alternative strategy, we expressed *N. lactamica* ATCC 23970 *ampG* (*ampG*NL23970) or *N. lactamica* ATCC 49142 *ampG* (*ampG*NL49142) in *N. gonorrhoeae* in lieu of the native gonococcal *ampG* (*ampG*GC). Expression of *N. lactamica ampG* in this background is driven by the native *ampG*GC promoter. Determination of the amount of PG monomers released by the allelic replacement mutants relative to the amount released by WT *N. gonorrhoeae* allows us to determine if *N. lactamica* AmpG is more, less, or equally efficient as AmpGGC. Expression of *ampG*NL23970, but not of *ampG*NL49142 in *N. gonorrhoeae* resulted in lower levels of PG monomer release compared to WT *N. gonorrhoeae* (Fig. 4). The gonococcal strain expressing *ampG*NL49142 released similar levels of PG monomer as WT *N. gonorrhoeae* (Fig. 4). These results suggest that AmpGNL49142 transports similar amounts of PG monomer as AmpGGC, while AmpGNL23970 is the most efficient at recycling of the three.

**Figure 4.**
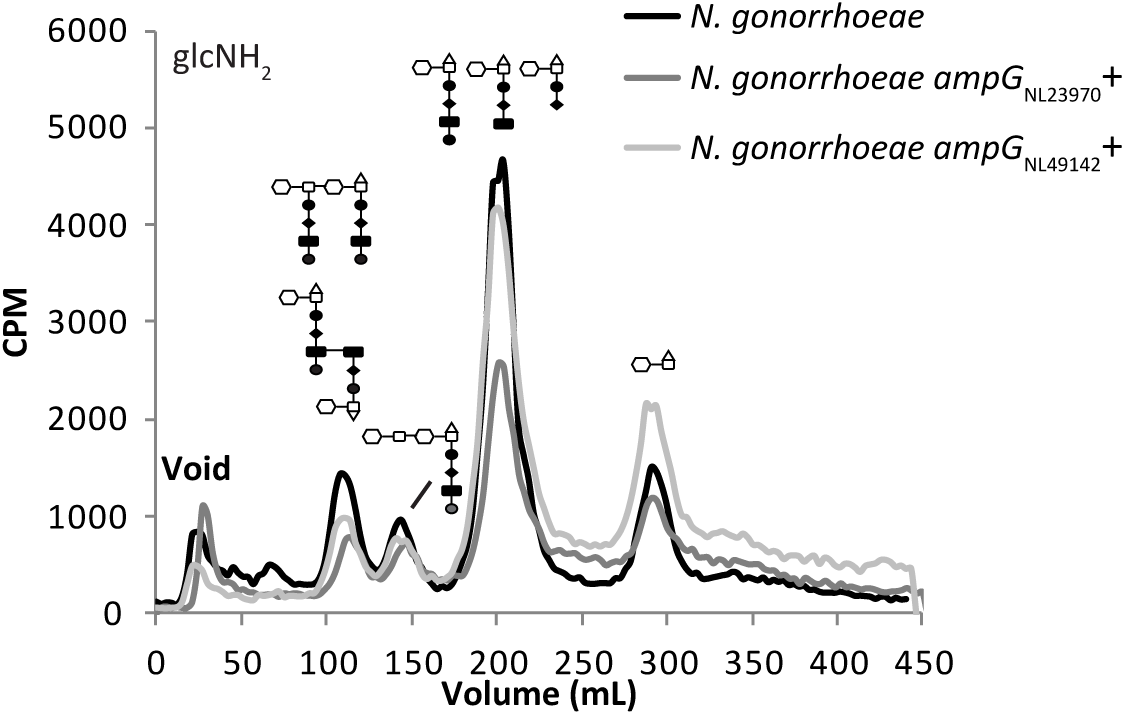
Gonococcal *ampG*^-^ strains expressing *ampG* from two different strains of *N. lactamica* released different amounts of PG monomer. The *ampG* gene from *N. lactamica* ATCC 49142 or *N. lactamica* ATCC 23970 were expressed in *N. gonorrhoeae* in lieu of the native gonococcal *ampG* gene. Expression of *ampG*NL23970 but not *ampG*NL49142 in *N. gonorrhoeae* reduced the amount of PG monomer released.

AmpGNL49142 and AmpGNL23970 differ by five residues. Previous work established that natural polymorphisms of gonococcal and meningococcal AmpG at residues 391, 398, and 402 contribute to differences in PG monomer release in these two pathogenic *Neisseria* [16]. Both isolates of *N. lactamica* code for the same amino acids at AmpG positions 391, 398, and 402. We compared AmpG sequences from the two *N. lactamica* isolates, *N. gonorrhoeae,* and *N. meningitidis*, and one of the five aforementioned residues is only found in AmpGNL49142. AmpG residue 272 is a threonine in *N. lactamica* ATCC 49142 and is an alanine in the other three species. We performed site-directed mutagenesis to determine if AmpG residue 272 plays a role in modulating AmpG efficiency in *N. lactamica*.

Expression of *ampG*NL49142T272A in *N. gonorrhoeae* lacking *ampG*GC resulted in lower levels of PG monomer release compared to WT *N. gonorrhoeae* (Fig. 5A), mimicking the phenotype seen when *ampG*NL23970 is expressed by *N. gonorrhoeae*. Additionally, expression of *ampG*NL23970A272T in *N. gonorrhoeae* resulted in WT-like levels of PG monomer release, which phenocopied the gonococcal strain expressing *ampG*NL49142 (Fig. 5B). We conclude that *N. lactamica* ATCC 23970 is better at recycling PG monomers compared to *N. lactamica* ATCC 49142 due to a polymorphism at AmpG site 272, which contributes in part to differences in the amount of PG fragments released by the two *N. lactamica* isolates.

**Figure 5.**
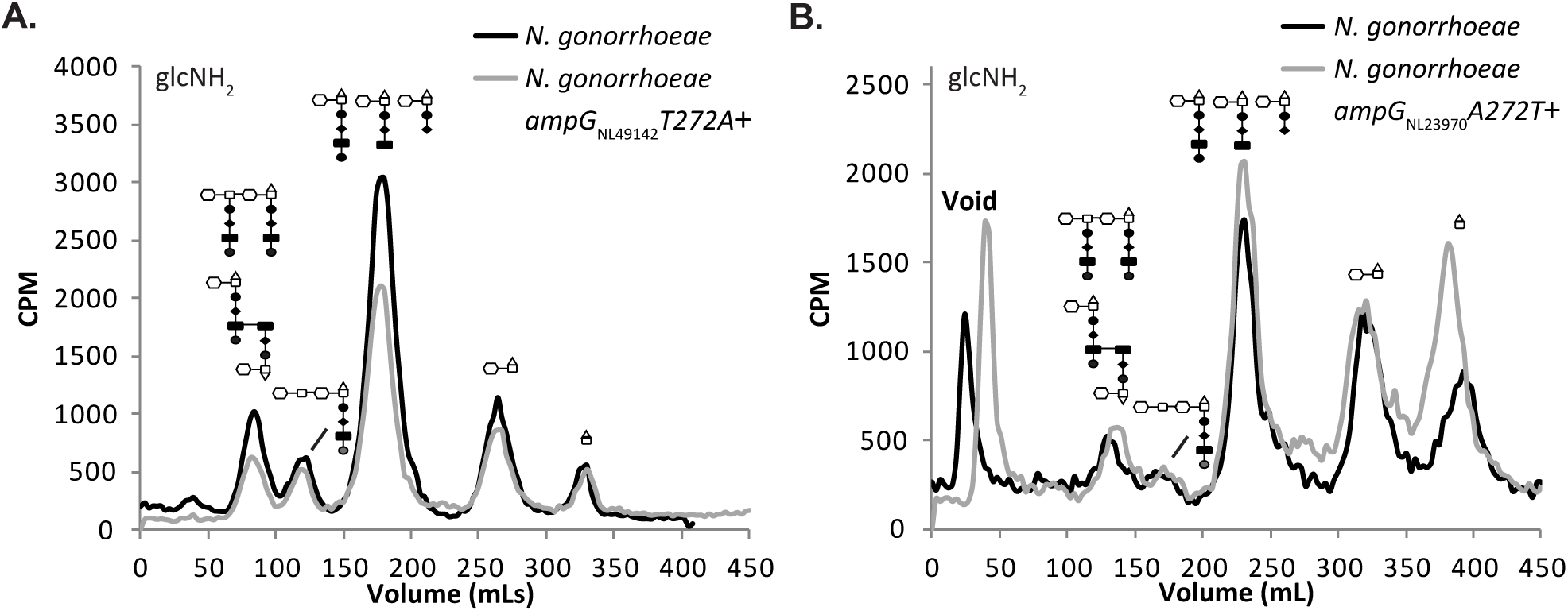
Polymorphism of *N. lactamica* AmpG contributes to the differences in the amounts of PG monomer released. A) *N. gonorrhoeae* lacking gonococcal *ampG* and instead expressing *ampG*NL49142 with a single substitution at AmpGNL49142 residue 272 from a threonine to an alanine shows reduced PG monomer release by approximately half. B) *N. gonorrhoeae* expressing *ampG*NL23970 with a single substitution at AmpGNL23970 residue 272 from an alanine to a threonine released *N. gonorrhoeae* WT- like levels of PG monomer.

We examined *N. lactamica* genome sequences to determine how common changes were at amino acid 272 of AmpG. Examination of 887 *N. lactamica* genome sequences found that 806 (91%) had alanine at residue 272, and 81 (9%) had threonine. No other amino acid was found at this position. Thus T272 is rare, predicting that most *N. lactamica* strains will have an AmpG that functions efficiently for PG fragment recycling.

### PG fragment release by animal-associated *Neisseria*

To determine if the macaque nasopharyngeal colonization model or the mouse oropharyngeal colonization model might be useful for examining the effects of PG fragment release in *Neisseria* infections, we characterized PG fragment release from macaque isolates AP312 and AP678 and from mouse isolate *N. musculi*. AP312 released PG monomers, free disaccharide, and anhydro-MurNAc, but little if any PG dimers (Fig. 6A). AP678 released PG dimers, PG monomers, and free disaccharide and monosaccharide (Fig. 6B). In addition, AP678 exhibited a large shoulder on the PG monomer peak, representing dipeptide PG monomers.

1. *N. musculi* released PG dimers, PG monomers, and free disaccharide (Fig. 7A). Examination of PG released using [2,6-^3^H]-DAP labeling demonstrated that *N. musculi* releases PG dimers, tetrasaccharide-peptide, PG monomers, and free peptides (Fig. 7B). In both the glucosamine labeling experiments and the DAP labeling experiments, PG monomers were the major PG fragments released.

**Figure 6.**
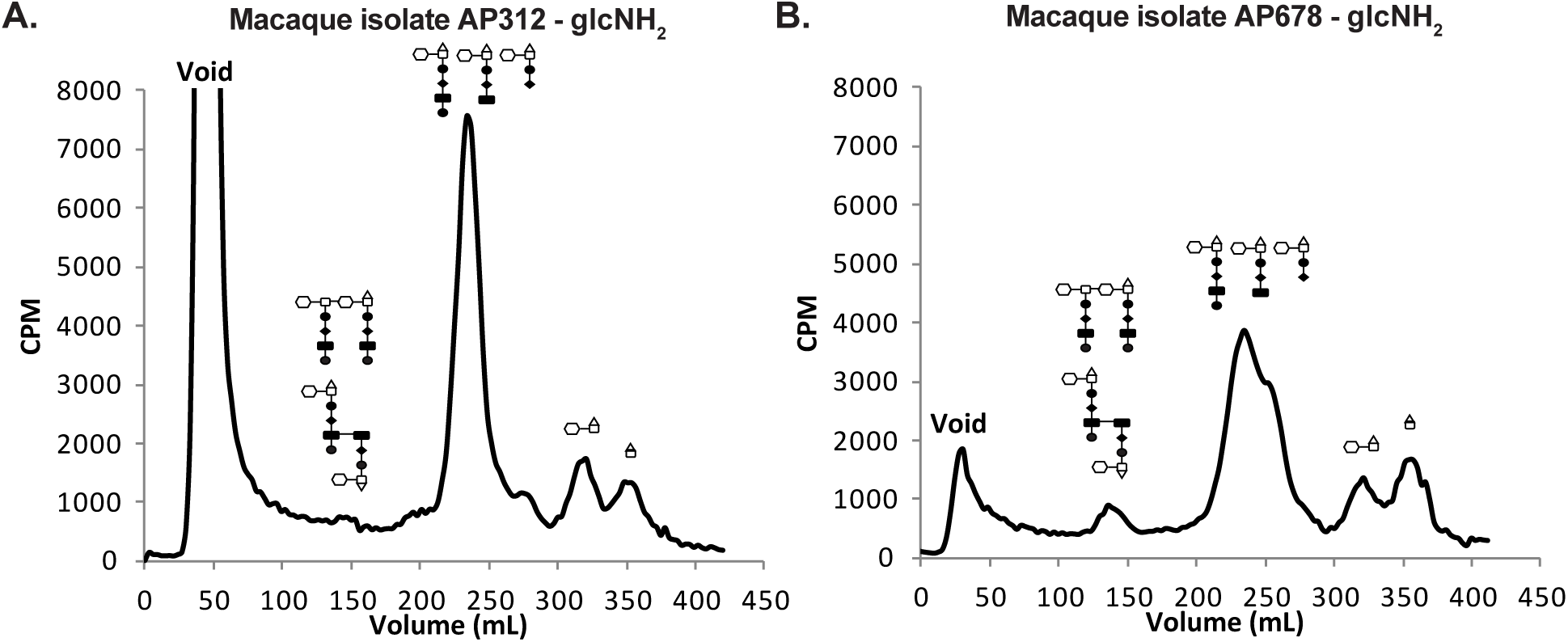
Macaque isolates AP312 and AP678 release different types and amounts of PG fragments.

**Figure 7.**
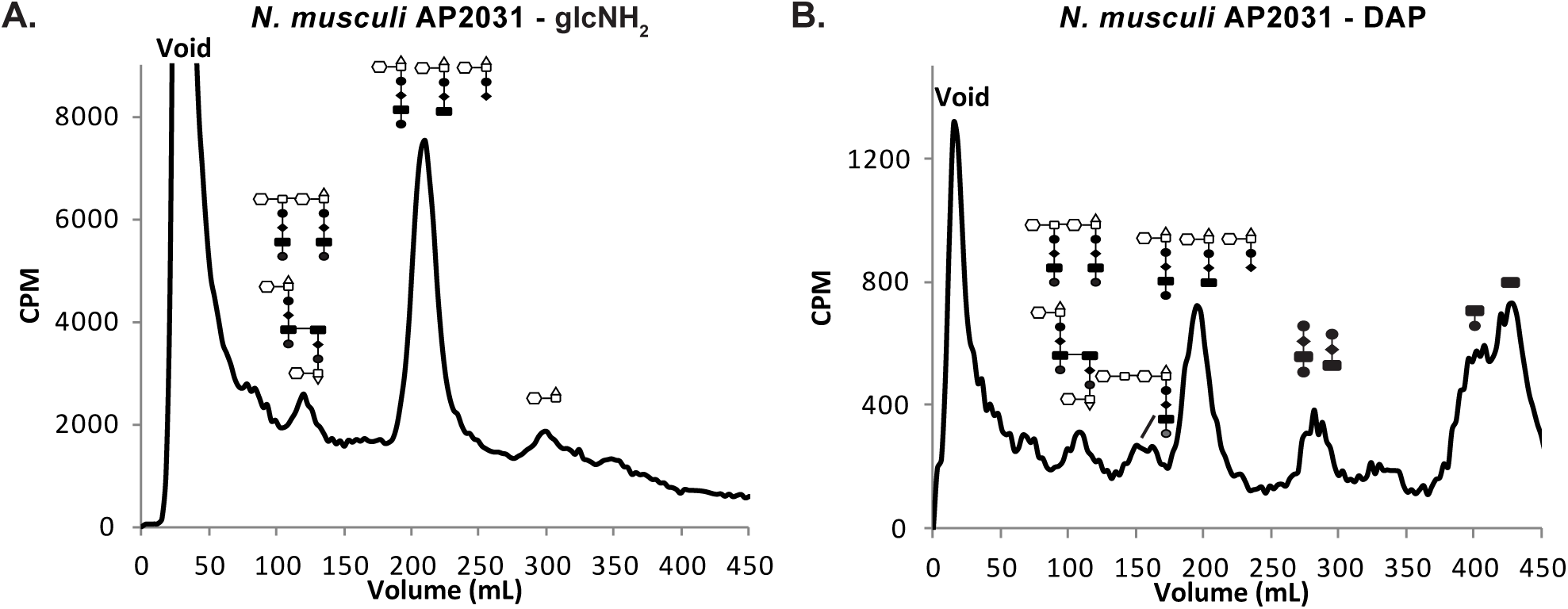
*N. musculi* releases a variety of PG fragments as determined form experiments using metabolic labeling with [^3^H]-glucosamine (A) or [^3^H]-DAP (B).

### Analysis of PG recycling by animal-associated *N. musculi* and macaque isolate AP312 using mutation of *ampG*

AmpG transports PG monomers and anhydro-disaccharide from the periplasm to the cytoplasm for recycling [34, 35]. By comparing the amount of PG released by *ampG* mutants compared to the amount released by WT strains, the normal amounts of glycan-containing PG fragments recycled and released can be calculated. We previously demonstrated that the human-associated species *N. mucosa* encodes a functional AmpG permease and recycles 95% of PG monomers liberated during growth [16].

We mutated *ampG* in *N. musculi* and macaque isolate AP312 by insertional inactivation with an antibiotic resistance marker and determined the PG fragment release profile of the *ampG* mutants. Mutation of *ampG* in *N. musculi* and macaque isolate AP312 resulted in an 8.7-fold and 10.8-fold increase in PG monomer release, respectively, with modest changes to anhydro-disaccharide release (Fig. 8A and 8B). *N. musculi* releases around 11% of PG monomer generated during growth, while macaque isolate AP312 releases approximately 9% of PG monomers. Like other *Neisseria* species, mutation of *ampG* did not alter PG dimer release in *N. musculi* or in macaque isolate AP312. This observation is consistent with the reported inability of *E. coli* AmpG to transport PG dimers and our previous results with *ampG* mutants of *N. gonorrhoeae* and *N. meningitidis* [19, 27, 35].

**Figure 8.**
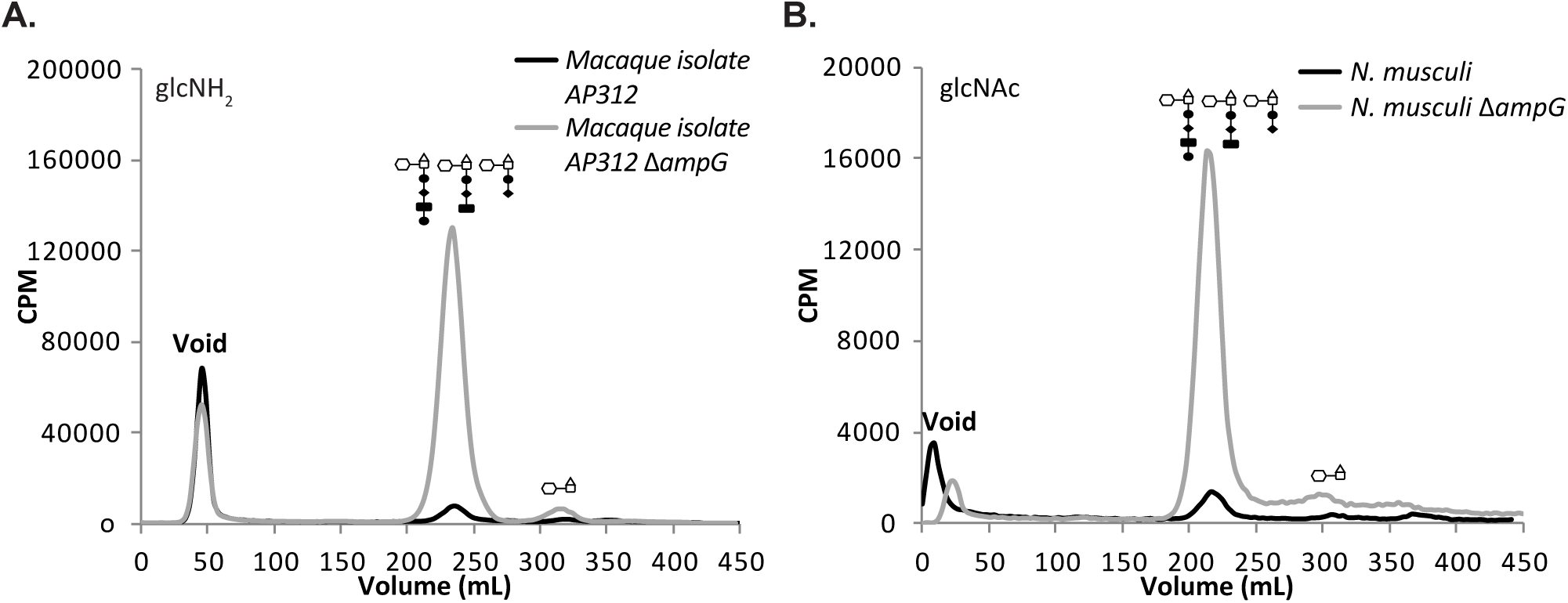
Mutation of *ampG* impaired PG fragment recycling in *N. musculi* and macaque symbiont AP312. Mutation of *ampG* in the macaque isolate AP312 (A) and *N. musculi* (B) resulted in large increases in the amount of PG monomer released, without affecting PG dimer release.

### PG fragments released by nonpathogenic human-associated *Neisseria* stimulate hNOD1 and hNOD2 activation in HEK293-reporter cells

Recognition of bacterial products by the immune system is important to prevent or resolve infections, as well as to attune the immune response [36, 37]. In humans, PG fragments are recognized by two intracellular receptors, human NOD1 (hNOD1) and NOD2 (hNOD2). hNOD1 responds to small PG fragments with a terminal DAP moiety [38] (Fig. 1B). hNOD2 binds to muramyl-dipeptide (MDP) and also responds to reducing-end PG monomers, which are generated from PG dimers or longer strands by the action of host lysozyme [23, 31, 39] (Fig. 1B). PG monomers released by *Neisseria* have anhydro-ends instead of reducing-ends due to the enzymatic activity of lytic transglycosylases that cleave the glycan backbone [28, 40, 41]. The NOD receptors signal through two major pathways to induce an immune response, and one of those pathways results in activation of NF-κB(48). We sought to determine if PG fragments released by nonpathogenic *Neisseria* are capable of stimulating an hNOD and an NF-κB dependent response.

To answer this question, we utilized HEK293 reporter cells that express a secreted alkaline phosphatase (SEAP) under the control of an NF-kB promoter and measured the amount of SEAP in the supernatant using a colorimetric assay. We treated the NOD1- or NOD2-expressing reporter cell lines with cell-free supernatant from *N. mucosa* or *N. lactamica* isolates, as well as supernatant from *N. gonorrhoeae* for comparison. Our initial hypothesis was that nonpathogenic *Neisseria* would induce lower levels of NOD activation. However, most of our nonpathogenic strains induced similar levels of hNOD1 and hNOD2 activation as *N. gonorrhoeae* (Fig. 9A and 9B). The value for *N. mucosa* stimulation of hNOD1 was lower than that for *N. gonorrhoeae*, though the values were not statistically different. A lower level of stimulation would be consistent with the decreased PG monomer release seen in *N. mucosa*. *N. lactamica* ATCC 23970 induced higher levels of hNOD2 activation than *N. gonorrhoeae* and other *Neisseria* species in this assay (Fig. 9B). The increased hNOD2 activation by *N. lactamica* ATCC 23970 could be due to PG dipeptide monomer, seen as a shoulder on the right side of the PG monomer peak, or to the substantial PG dimer release from *N. lactamica* ATCC 23970 (Fig. 3A).

**Figure 9.**
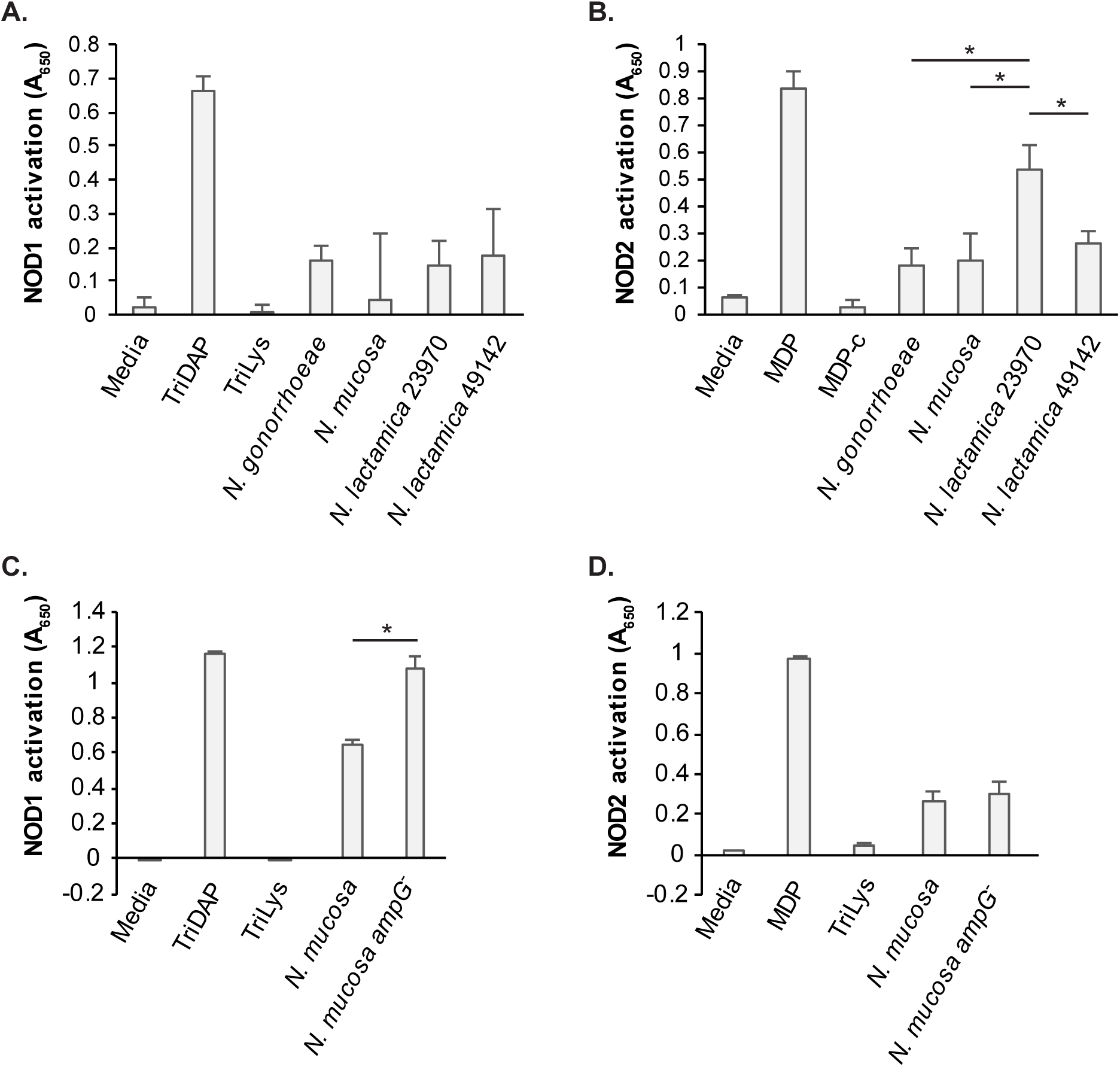
hNOD1 and hNOD2 activation by supernatants from different *Neisseria* species. Supernatants from *N. mucosa*, *N. lactamica* (A,B) and *ampG* mutants of and *N. mucosa* (C,D) induced hNOD1 (A,C) and hNOD2 (B,D) activation in HEK293 reporter cells overexpressing NOD1 or NOD2. Supernatant from an *ampG* mutant of *N. mucosa* induced a larger hNOD1 response compared to wild-type. Supernatant from *N. lactamica* ATCC23970 induced the largest hNOD2 response of all supernatant treatment samples. Statistical significance was determined using Student’s t-test. * indicates p<0.05.

Since mutation of *ampG* increases the amount of PG monomers released, and tripeptide monomers are known agonists of hNOD1, we hypothesized that supernatant from *ampG* mutants of *N. mucosa* would induce higher levels of hNOD1 activation compared to WT. As expected, supernatant from the *ampG* mutant activated hNOD1, but not hNOD2, to a greater degree compared to that of the WT parent (Fig. 9C and 9D). Thus, we conclude that *N. mucosa* and *N. lactamica* release hNOD- activating PG fragments, and that PG recycling modulates the amount of hNOD1 agonists released by the bacteria.

### PG fragments released by *N. musculi* stimulate mNOD1 activation in HEK293-reporter cells

Similar to hNOD1 expressed in humans, mice also express a NOD1 receptor. While hNOD1 is activated by tripeptide monomer and tripeptides, mNOD1 is activated by tetrapeptide monomer and tetrapeptides [32]. mNOD1 activation in HEK293-reporter cells was analyzed for *ampG* and *ldcA* mutant *N. musculi* strains (Fig. 10). Mutation of *ampG* in *N. musculi* resulted in mNOD1 activation slightly higher than seen with the WT, though this apparent difference was not statistically significant. We mutated *ldcA* in *N. musculi* since LdcA is the enzyme which converts tetrapeptides into tripeptides in the periplasm [42]. Since free tetrapeptides and tetrapeptide monomers are the mNOD1 agonists, and tripeptides versions of those molecules are poor stimulatory molecules for mNOD1, the mutation would make all the released PG monomers and free peptides into mNOD1 agonists. Supernatant from the *ldcA* mutant stimulated mNOD1 to a much higher degree than WT or *ampG* supernatants. We created a genetic construct for expression of genes of interest at a neutral site on the *N. musculi* chromosome, and we used this construct to overexpress *ldcA*. Overexpression of *ldcA* in *N. musculi* resulted in a background level of mNOD1 stimulation (Fig. 10).

**Figure 10.**
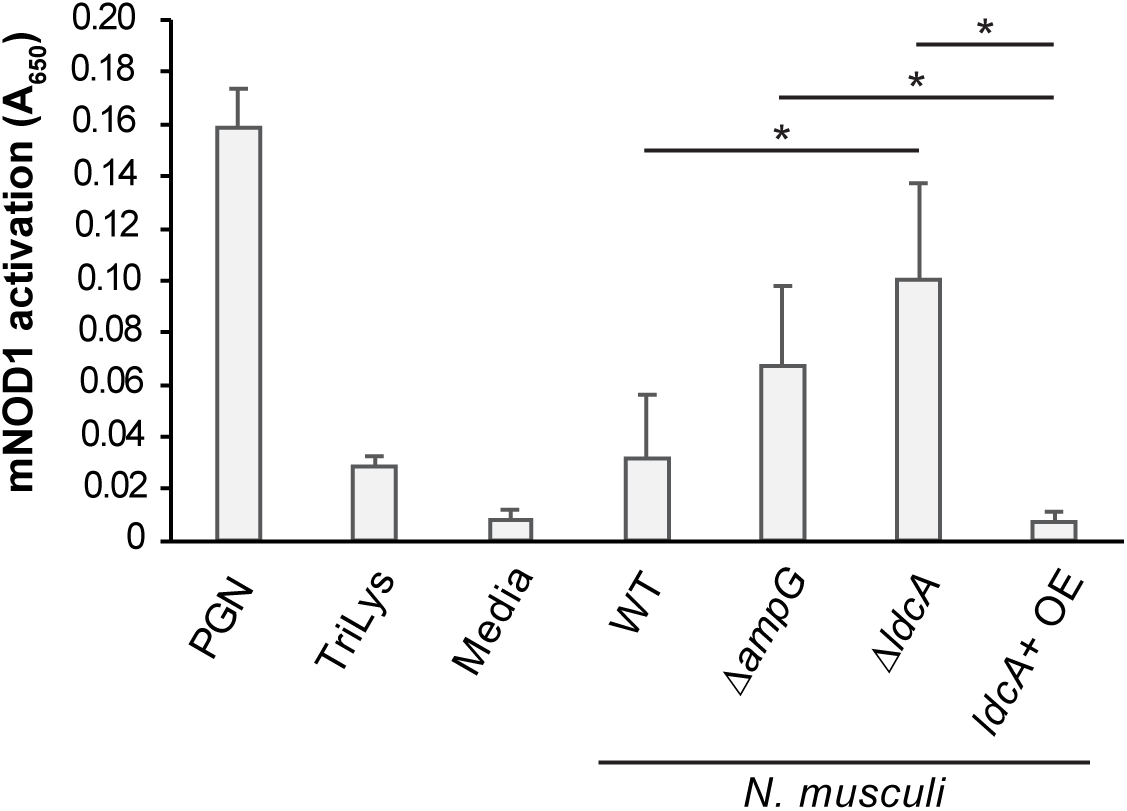
mNOD1 activation by supernatants from *N. musculi* species used to treat HEK-293 cells overexpressing mNOD1. Supernatants from *N. musculi* WT, *ampG* or *ldcA* mutants, or a strain overexpressing *ldcA* were used. Statistical significance was determined using Student’s t-test. * indicates p<0.05.

## Discussion

Due to the role of PG monomers in causing damage to mucosal epithelium by inducing death and sloughing of ciliated cells during *N. gonorrhoeae* and *Bordetella pertussis* infections, we hypothesized that nonpathogenic *Neisseria* would release only small amounts of inflammatory PG fragments. The first characterization of PG fragments released by *N. mucosa*, performed as quantitative fragment release in comparison to *N. gonorrhoeae*, showed that this species releases lower levels of proinflammatory PG monomers compared to *N. gonorrhoeae* and no PG dimers [16]. It was logical to infer that the reduced PG monomer release by *N. mucosa* means that nonpathogenic *Neisseria* are less inflammatory compared to *N. gonorrhoeae*. This inference is supported by the observation that human-associated, nonpathogenic *Neisseria* induce lower Toll-like receptor (TLR4) response compared to *N. gonorrhoeae* and *N. meningitidis* due to differences in lipooligosaccharide modification [43].

As we continued to investigate PG fragment release by more strains of human- and animal-associated *Neisseria*, it became clear that the situation is not as simple as we initially hypothesized. There are both interspecies and intraspecies variations in PG fragment release patterns. *N. mucosa* and macaque isolate AP312 did not release PG dimers, while *N. lactamica*, *N. musculi*, and macaque isolate AP678 released PG dimers. The two strains of *N. lactamica* evaluated showed differences in the relative amounts of PG monomer and PG peptides released (Fig. 3). Labeling of *N. mucosa* with [^3^H]- DAP revealed that although *N. mucosa* released less PG monomer compared to *N. gonorrhoeae*, *N. mucosa* released more PG-derived peptides that might also activate hNOD1 (Fig. 2). *N. meningitidis*, which despite its ability to cause invasive disease is more commonly found as a colonizer of the nasopharyngeal space, also releases less PG monomer and more PG peptides compared to *N. gonorrhoeae* [19, 30].

The two strains of *N. lactamica* used in the study displayed differences in the relative amounts of PG monomer and PG peptides released, partly due to differences in the efficiency of PG recycling. Such intraspecies variation in PG monomer release has been previously observed with two different isolates of *N. meningitidis* that have polymorphisms at AmpG residues 398 and 402 [16]. We identified another AmpG residue (residue 272) capable of modulating PG recycling efficiency and indirectly controlling the amount of PG monomer released. AmpG residue 272 is a threonine in *N. lactamica* ATCC 49142 and an alanine in *N. lactamica* ATCC 23970, and WT *N. gonorrhoeae* and WT *N. meningitidis*. This finding suggests that different species of *Neisseria* independently evolved strategies to fine-tune the amount of PG fragments released. Examination of 887 *N. lactamica* sequences available in public databases found that T272 in AmpG is present in 9% of strains. So, only a minority of *N. lactamica* strains are predicted to release larger amounts of PG fragments. Information about carrier state or disease in the people providing these isolates was not available, and thus it is not known if the amino acid substitution is more associated with disease isolates. It is possible that alterations to PG fragment release would allow strains to colonize different body sites more effectively.

Through hNOD1and hNOD2 activation we were able to determine that despite the nonpathogenic nature of human-associated *Neisseria* commensals, NOD activation was not lower in these species compared with *N. gonorrhoeae.* In fact, hNOD2 activation was higher in *N. lactamica* ATCC 23970 than in the pathogenic *N. gonorrhoeae.* Therefore, PG fragment release in significant amounts does not make commensal *Neisseria* pathogens during nasopharyngeal colonization. Further research into the differences between NOD responses in the different niches infected by *Neisseria* may lead to a better understanding of these responses.

The recent establishment of macaque and mice-associated *Neisseria* as model organisms provide avenues for studying aspects of *Neisseria* colonization and infection in their cognate host organisms [6, 7, 13]. One caveat of using mice as a model organism for understanding PG fragment responses in humans is that the murine NOD1 detects tetrapeptide monomers instead of tripeptide monomers like the human NOD1 [32]. Still, our findings indicate that both macaque isolates and *N. musculi* release PG monomers, and that macaque isolate AP312 and *N. musculi* recycle PG fragments liberated during turnover (Fig. 6 and 7). Our ability to mutate genes in macaque isolate AP312 and *N. musculi* provide us with tools to vary the amounts and types of PG fragments that the host organism is exposed to, and to study the corresponding host response. Future studies in these models will allow us to better understand the roles of PG fragments in *Neisseria* infection and discern factors that distinguish a pathogen from a nonpathogen.

